# Intrinsic noise improves speech recognition in a computational model of the auditory pathway

**DOI:** 10.1101/2020.03.16.993725

**Authors:** Achim Schilling, Richard Gerum, Alexandra Zankl, Claus Metzner, Andreas Maier, Patrick Krauss

## Abstract

Noise is generally considered to harm information processing performance. However, in the context of stochastic resonance, noise has been shown to improve signal detection of weak subthreshold signals, and it has been proposed that the brain might actively exploit this phenomenon. Especially within the auditory system, recent studies suggest that intrinsic noise plays a key role in signal processing and might even correspond to increased spontaneous neuronal firing rates observed in early processing stages of the auditory brain stem and cortex after hearing loss. Here we present a computational model of the auditory pathway based on a deep neural network, trained on speech recognition. We simulate different levels of hearing loss and investigate the effect of intrinsic noise. Remarkably, speech recognition after hearing loss actually improves with additional intrinsic noise. This surprising result indicates that intrinsic noise might not only play a crucial role in human auditory processing, but might even be beneficial for contemporary machine learning approaches.

## Introduction

The term *noise* usually describes undesirable disturbances or fluctuations, and is considered to be the *“fundamental enemy”* [1] for communication and error-free information transmission and processing in engineering. However, a vast and still increasing number of publications demonstrate the various benefits of noise for signal detection and processing, among which the most important phenomena are called stochastic resonance [1], coherence resonance [2], and recurrence resonance [3].

The term stochastic resonance (SR), first introduced by Benzi et al. [4], refers to a processing principle in which signals that would otherwise be sub-threshold for a given sensor can be detected by adding a random signal of appropriate intensity to the sensor input [4–6]. SR occurs ubiquitously in nature and covers a broad spectrum of systems in physical and biological contexts [1, 7]. Especially in neuroscience, it has been demonstrated to play an essential role in a vast number of different systems [8–16]. Also, it has already been proposed that spontaneous random activity, i.e. noise, may increase information transmission via SR in the auditory brain stem [17].

In self-adaptive signal detection systems based on SR, the optimal noise intensity is continuously adjusted via a feedback loop so that the system response remains optimal in terms of information throughput, even if the characteristics and statistics of the input signal change. The term adaptive SR was coined for this processing principle [18–20]. In a previous study we demonstrated that the auto-correlation of the sensor output, a quantity always accessible and easy to analyze by neural networks, can be used to quantify and hence maximize information transmission even for unknown and variable input signals [21].

In further studies we demonstrated theoretically and empirically that adaptive SR based on output auto-correlations might be a major processing principle of the auditory system that serves to partially compensate for acute or chronic hearing loss, e.g. due to cochlear damage [22–25]. Here, the noise required for SR would correspond to increased spontaneous neuronal firing rates in early processing stages of the auditory brain stem and cortex, and would be perceived as a phantom perception. Remarkably, this phenomenon has frequently been observed in animal models and in humans with subjective tinnitus [26–29], which in turn is assumed to be virtually always caused by some kind of apparent [30–33] or hidden hearing loss [34, 35]. From this point of view, phantom perceptions like tinnitus seem to be a side effect of an adaptive mechanism within the auditory system whose primary purpose is to compensate for reduced input through continuous optimization of information transmission [22, 24, 25].

The dorsal cochlear nucleus (DCN) was shown to be the earliest processing stage where acoustic trauma causes increased spontaneous firing rates [29, 36–38]. Interestingly, this increase in spontaneous activity, i.e. neural hyperactivity, is correlated with the strength of the behavioral signs of tinnitus in animal models [39, 40]. Furthermore, the hyperactivity is localized in those regions of the DCN that are innervated by the damaged parts of the cochlear [41]. Gao and colleagues [42] recently described changes in DCN fusiform cell spontaneous activity after noise exposure that supports the proposed SR mechanism. In particular, the time course of spontaneous rate changes shows an almost complete loss of spontaneous activity immediately after loud sound exposure (as no SR is needed due to stimulation that is well above threshold), followed by an overcompensation of spontaneous rates to levels well above pre-exposition rates since SR is now needed to compensate for acute hearing loss [42]. It is well known that the DCN receives not only auditory input from the cochlea, but also from the somatosensory system [29, 43, 44], and that noise trauma alters long-term somatosensory-auditory processing in the DCN [45], i.e. somatosensory projections are up-regulated after deafness [46].

Therefore, we previously proposed the possibility that the neural noise for SR is injected into the auditory system via somatosensory projections to the DCN [22, 24, 25]. The idea that SR plays a key role in auditory processing and actually takes place in the DCN is supported by a number of findings. For instance, it is well known, that jaw movements lead to a modulation of subjective tinnitus loudness [47]. This may easily be explained within our framework, as jaw movements alter somatosensory input to the DCN. Since this somatosensory input corresponds to the noise required for SR, auditory input to the DCN is modulated through this mechanism, and the altered noise level is then perceived as modulated tinnitus [22, 24, 25]. Along the same line, one may explain why both, the temporo-mandibular joint syndrome and whiplash, frequently cause so called somatic tinnitus [48]. Another example is the finding of Tang et al. [49, 50], who demonstrated that somatosensory input and hence tinnitus sensation may also be modified by serotonergic regulation of excitability of principal cells in the DCN. In addition, DCN responses to somatosensory stimulation are enhanced after noise-induced hearing loss [51,52]. Finally, and most remarkable, electro-tactile stimulation of finger tips, i.e. increased somatosensory input, significantly improves both, melody recognition [53] and speech recognition [54] in patients with cochlear implants.

In order to further support the hypothesis that SR plays a key role in auditory processing and takes place in the DCN, we here present a hybrid computational model of the auditory pathway, trained on speech recognition. In particular, we simulate different levels of hearing loss (cochlear damage) and compare the resulting accuracies for speech recognition with the accuracy of the non-disturbed model (i.e. without simulated hearing loss). As expected, we find that the accuracy decreases systematically with increasing hearing loss.

Subsequently, we add intrinsic noise of different intensities to the model. Here, we find SR-like behavior for all levels of hearing loss: depending on the intensity of the noise, accuracy first increases, reaches a peak, and finally decreases again. This means that speech recognition after hearing loss may indeed be improved by our proposed mechanism. This intriguing result indicates, that SR indeed plays a crucial role in auditory processing, and might even be beneficial for contemporary machine learning approaches.

## Results

### Layout of the computational model and general approach

The model comprises three modules (Figure 1): (1) an artificial cochlea modeled as an array of band-pass filters, (2) a model of the dorsal cochlear nucleus (DCN), implemented as an array of leaky integrate-and-fire (LIF) neurons, and (3) a deep neural network, that represents all further processing stages beyond the DCN up to the auditory cortex and higher, language associated, cortex areas.

**Figure 1:**
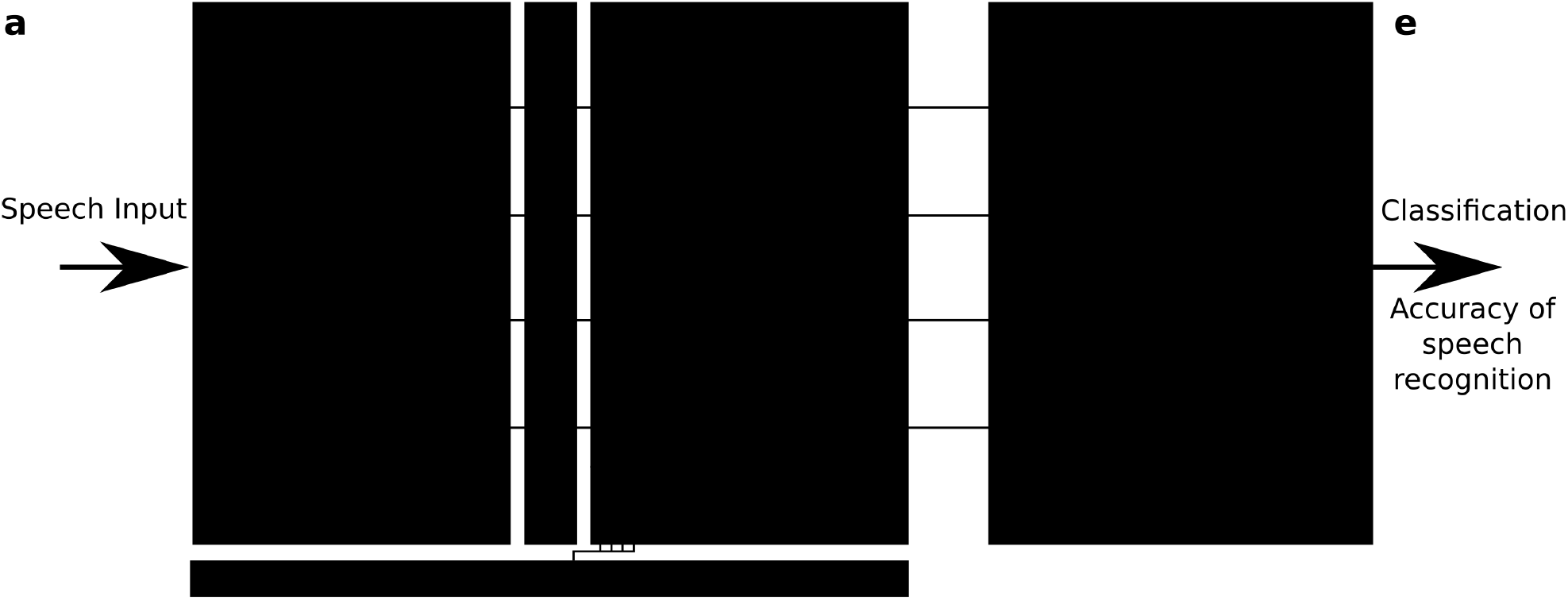
Model layout. The complete model consists of three different modules representing different stages of the auditory pathway in the human brain. The input to the model are single words encoded as wave files with a sampling rate of 44.1 kHz and 1s duration (a). The cochlea and the spiral ganglion are modeled as an array of 30 band-pass filters (b). The continuous output signal of (b) serves as input to 30 leaky integrate-and-fire-neurons representing the DCN (c). The spike-train output of the DCN model is down sampled and serves as input for a deep neural network that is trained with error backpropagation on the classification of 207 different german words (d). The classification accuracy serves as a proxy for speech recognition (e). In order to investigate the effect of a particular hearing loss, the cochlea output amplitude is decreased by a certain factor independently for all frequency channels (f). White noise representing somatosensory input to the DCN can be added independently to the input of the different leaky-integrate-and-fire-neurons (LIF, g).

The input to the model are single words of spoken language encoded as wave files with a sampling rate of 44.1 kHz and 1 s duration (Figure 1a, cf. Methods). These wave files represent the acoustic input of speech to the auditory system, and are processed in the first module of the model representing the cochlea and the spiral ganglion (Figure 1b). Like in previously published models [55–57], this module is implemented as an array of rectangular band-pass filters. In order to limit the total computation time, we restricted our model to 30 band-pass filters, instead of the actual amount of approximately 3500 inner hair cells in the human cochlea [58]. According to the physiology of the cochlea [59], the center frequencies of the band-pass filters are chosen such that they cover the frequency range from 100 Hz to 10 kHz in logarithmic steps (cf. Methods).

The continuous multi-channel output of the band-pass filter array serves as input to an array of 30 LIF neurons [60] representing the DCN (Figure 1c). We here applied a one-to-one mapping from band-pass filters to model neurons, i.e. we do not explicitly account for putative cross-talk between neighboring frequency channels. However, since both the cochlea and the DCN model only consist of 30 different frequency channels, each of these channels may be regarded as an already coarse grained version of approximately 100 different frequency channels that exist in the human auditory system. Thus, eventual cross-talk is implicitly implemented in our model within each of the 30 modelled channels. The output of our DCN model comprises the spike trains of the 30 LIF neurons. Note that, in our DCN model, a single IF neuron represents approximately ten biological neurons processing the same frequency channel [61].

In our cochlea and DCN model, the outputs of the band-pass filters and the membrane potentials of the LIF neurons change with the same rate (44.1 kHz) as the wave file input. However, the LIF neurons spike at lower average rates, due to their refractory period. It is therefore possible to downsample this sparse output spike train, thereby reducing the data volume for the subsequent deep neural network. In order to preserve enough temporal information for phase coding, we down-sample the DCN output only by a factor of five, so that the 44100 momentary amplitudes of the input wave file per second are finally transformed into a binary 30 × 8820 matrix.

These binary matrices serve as training input for the deep neural network, representing all further processing stages beyond the DCN up to the auditory cortex and higher, language associated, cortex areas. The neural network consists of four convolutional layers and three fully connected layers, and is trained with error backpropagation on the classification of 207 different German words (Figure 1d). The resulting classification accuracy of the trained network serves as a proxy for speech recognition (Figure 1e).

In order to simulate a particular hearing loss, the output amplitudes of the cochlea model are decreased by a certain factor, independently for the different frequency channels (Figure 1f). Subsequently, these modified cochlea outputs are further processed in the LIF neurons, finally resulting in a new binary matrix for each word for a particular hearing loss. These new matrices then serve as test data for the previously trained deep neural network, yielding a new classification accuracy. By comparing the reference test accuracy (without any hearing loss) with the new test accuracy, the effect of a particular hearing loss on speech recognition was estimated.

Optionally, Gaussian noise with zero mean and a certain standard deviation, representing somatosen-sory input to the DCN, was added independently to the input of each LIF neuron (Figure 1g). Here, the standard deviation corresponds to the noise intensity. As described before, again this finally results in a new binary matrix for each wave file, yet corresponding to a particular hearing loss and, in addition, also to a particular set of frequency channel specific noise intensities. Again, all these new matrices serve as test data for the deep neural network. By comparing the reference test accuracy (without any hearing loss and noise) with the new test accuracy, the effect of particular noise intensities on speech recognition with a certain hearing loss was estimated. A sketch of the complete data flow in case of certain hearing loss and additional noise is depicted in Figure 2.

**Figure 2:**
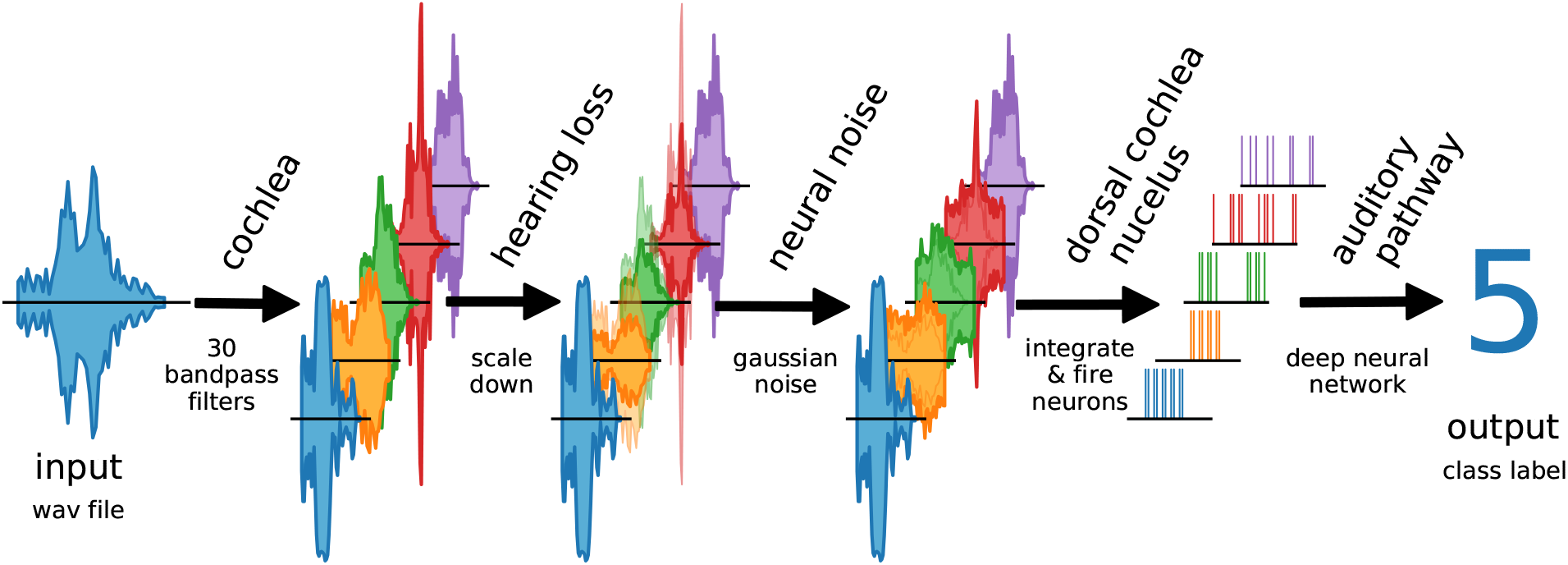
Data flow in auditory pathway model. The scheme shows how the speech data is processed within the model. The cochlea splits the signal via 30 bandpass filters. The bandpass filtered data is scaled down to simulate a hearing loss. The hearing loss affects only channels within the speech relevant frequency range (orange, green, red). The other frequency channels are unchanged. Neural noise is added to investigate the effect of stochastic resonance (only in hearing impaired channels). The DCN is simulated as 30 LIF neurons. Each LIF neuron represents a complete biological neuron population. The spike data is down-sampled and fed to the deep neural network.

### DCN model neurons show phase coupling below 4 kHz

In order to validate our DCN model, we investigate the spike train output of the 30 LIF neurons for different sine wave inputs (Figure 3). As described in the Methods section, the parameters of the LIF neurons are chosen so that the refractory time (0.25 ms) of the neurons does not allow for firing rates above 4 kHz. Thus, one LIF neuron in our model represents approximately ten biological neurons, having individual refractory times above 1 ms.

**Figure 3:**
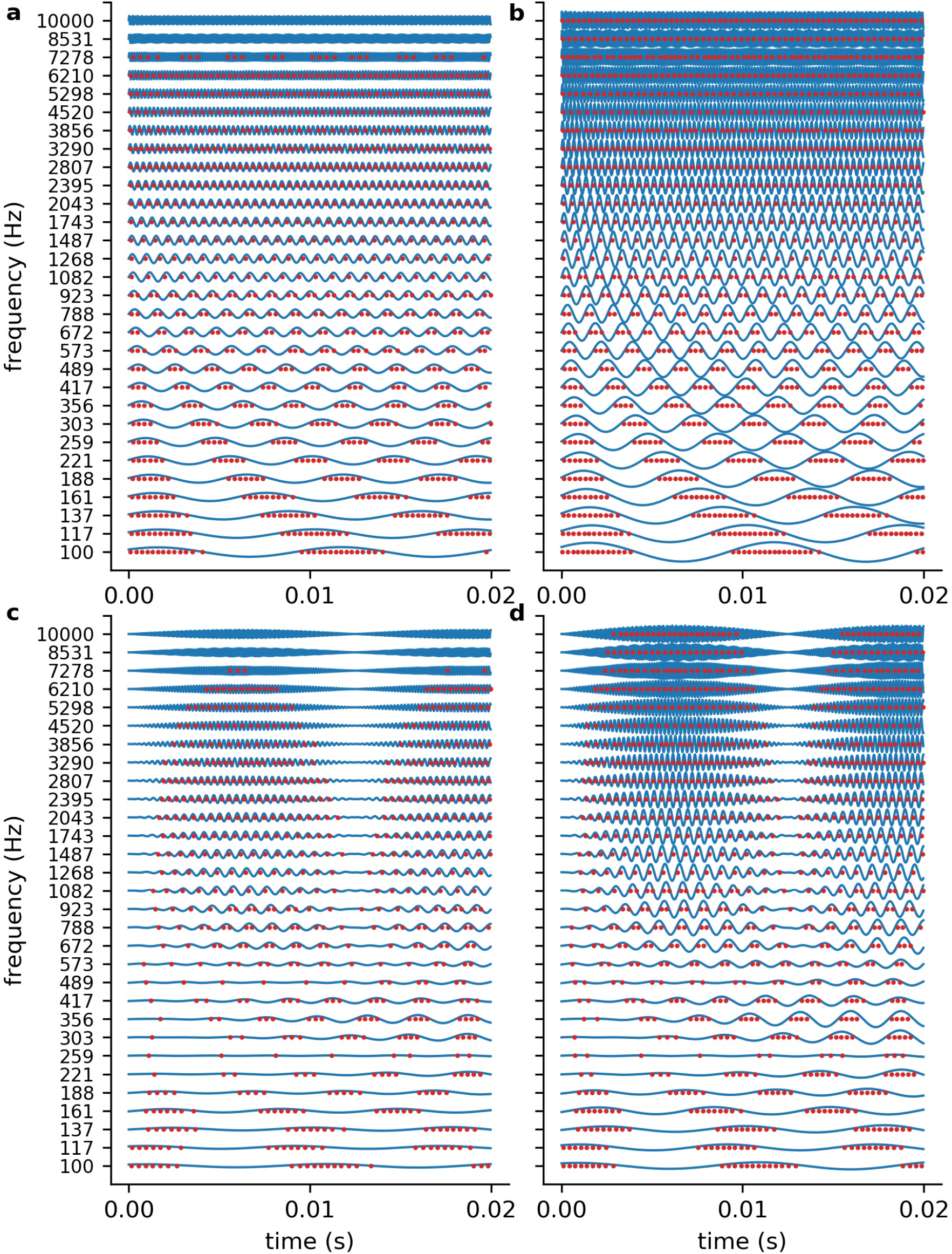
DCN model response to sine waves. Shown are the spiking outputs of the LIF neurons for sine input with two different constant amplitudes (a: 0.001, b: 0.002), and two different amplitude modulations (c, d). For lower amplitudes (a) and higher frequencies the LIF neurons do not spike at all, whereas for higher amplitudes a rate code can be observed as the neurons’ maximum spiking rate is limited due to the refractory period. The parameters of the LIF neurons are chosen so that there is phase coupling in the frequency range which is relevant for speech perception.

We find that for stimulus frequencies above 4 kHz and amplitudes of 0.001 the LIF neurons do not spike at all (Figure 3a). In contrast, for a larger amplitude of 0.002, a rate coding without phase coupling can be observed (Figure 3b). Furthermore, we find that the LIF neurons are sensitive to amplitude modulations also in the frequency range above 4 kHz (Figure 3c, d). Thus, our DCN neurons are designed so that they allow for phase coupling in the frequency range crucial for speech comprehension, as is known from the human auditory system.

### Word processing from cochlea to DCN

In analogy to the auditory system, the complex auditory stimuli representing spoken words (Figures 4a and 2) are transformed in the cochlea into continuous signals in a number of different frequency channels, in our model 30. However, the cochlea does not perform a simple Fourier transform, but rather splits the signal into multiple band pass filtered signals, thereby preserving the complete phase information (Figures 4b and 2). The auditory nerve fibers directly transmit this analogue signal to the DCN [62], which is a special feature of the auditory system [61].

**Figure 4:**
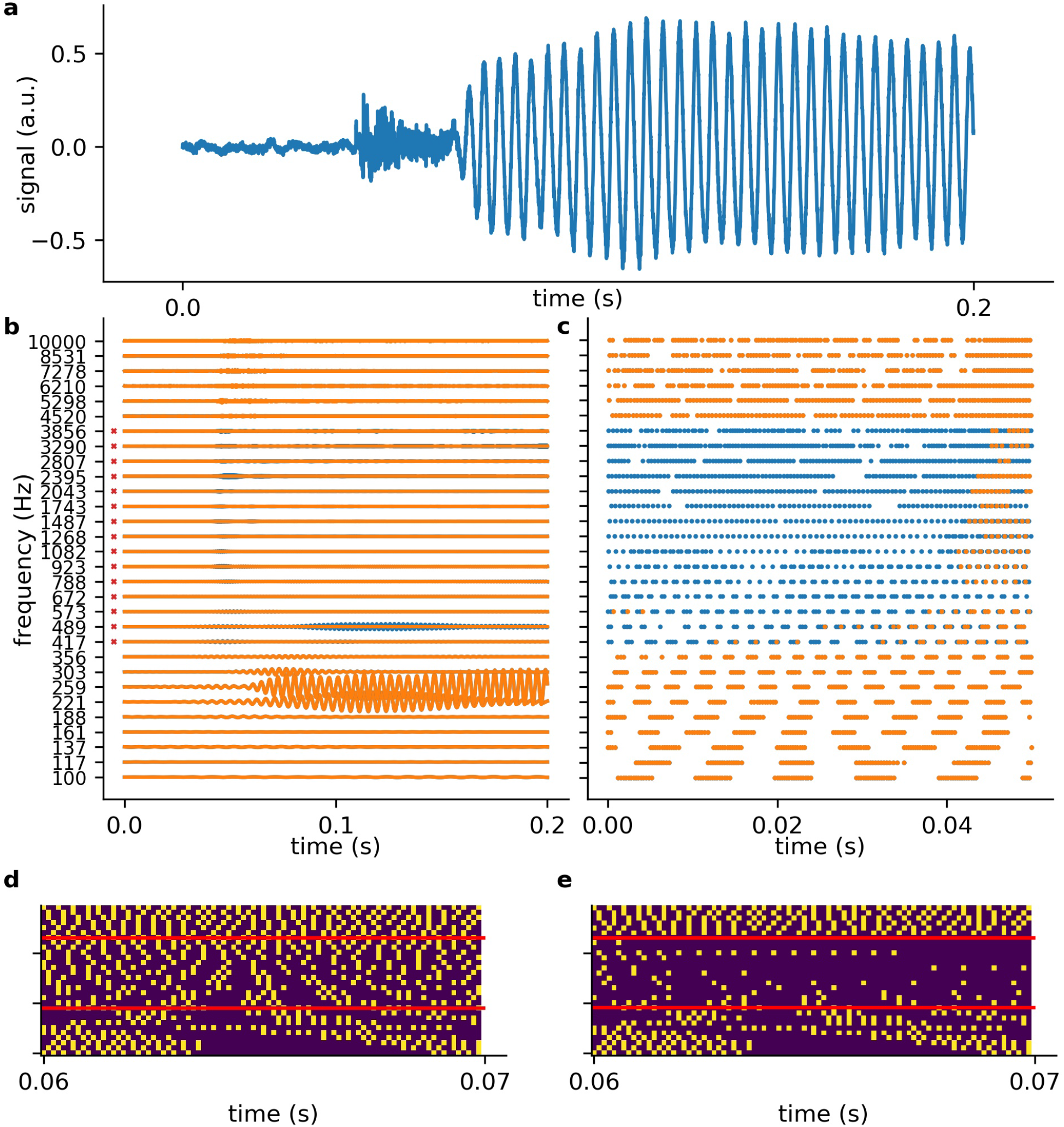
Exemplary processing of a word in cochlea and DCN model. a: The first 0.2s of audio data of the German word “die” (the). b) The 30 frequency components (blue without hearing loss, orange with hearing loss) after the first part of the model, which represents the cochlea and the spiral-ganglion (Fig. 1a). A virtual hearing loss is applied by weakening the signal at a certain frequency range (e.g. 400 Hz–4kHz, −30 dB). The bandpass filtered signal (matrix of 30 frequency channels and *f_s_* × signal duration) is fed to the LIF neurons (refractory time: ≈ 0.25*ms*) and spike trains (c) are generated. These spike trains are down-sampled by a factor of 5 and fed to the deep neural network (d). e: The same signal (of d) with added hearing loss of 30dB in the frequency range 400 Hz–4 kHz being the speech relevant range.

The analogue signals are then further transformed into spike train patterns in the DCN (Figures 2 and 4c). Thus, each spoken word is represented as a unique spiking pattern with a dimensionality of 30 × *N*, where 30 corresponds to the number of frequency channels and *N* is the sampling rate in Hz times the word length in seconds. Note that we down-sampled these matrices by a factor of five from 44100 Hz to 8200 Hz for deep learning (Figure 4d). This does not affect the phase coupling information in the speech relevant frequency range. In order to analyze speech processing in an impaired auditory system, we simulated a hearing loss in the speech relevant frequency range (400 Hz–4 kHz) by decreasing the cochlea output amplitudes by a certain factor. The weakened cochlea outputs and the resulting modified DCN spike train outputs are shown in Figure 4b and c, where orange corresponds to an exemplary hearing loss of 30 dB, and blue corresponds to the undisturbed signals, i.e. without hearing loss. The corresponding down sampled spike pattern matrices used as test data for the deep neural network, are shown in Figure 4d (without hearing loss) and in Figure 4e (with 30 dB hearing loss). We provide an exemplary overview of the effect of different hearing losses from 0 dB to 45 dB on the spike pattern matrices in Figure 5.

**Figure 5:**
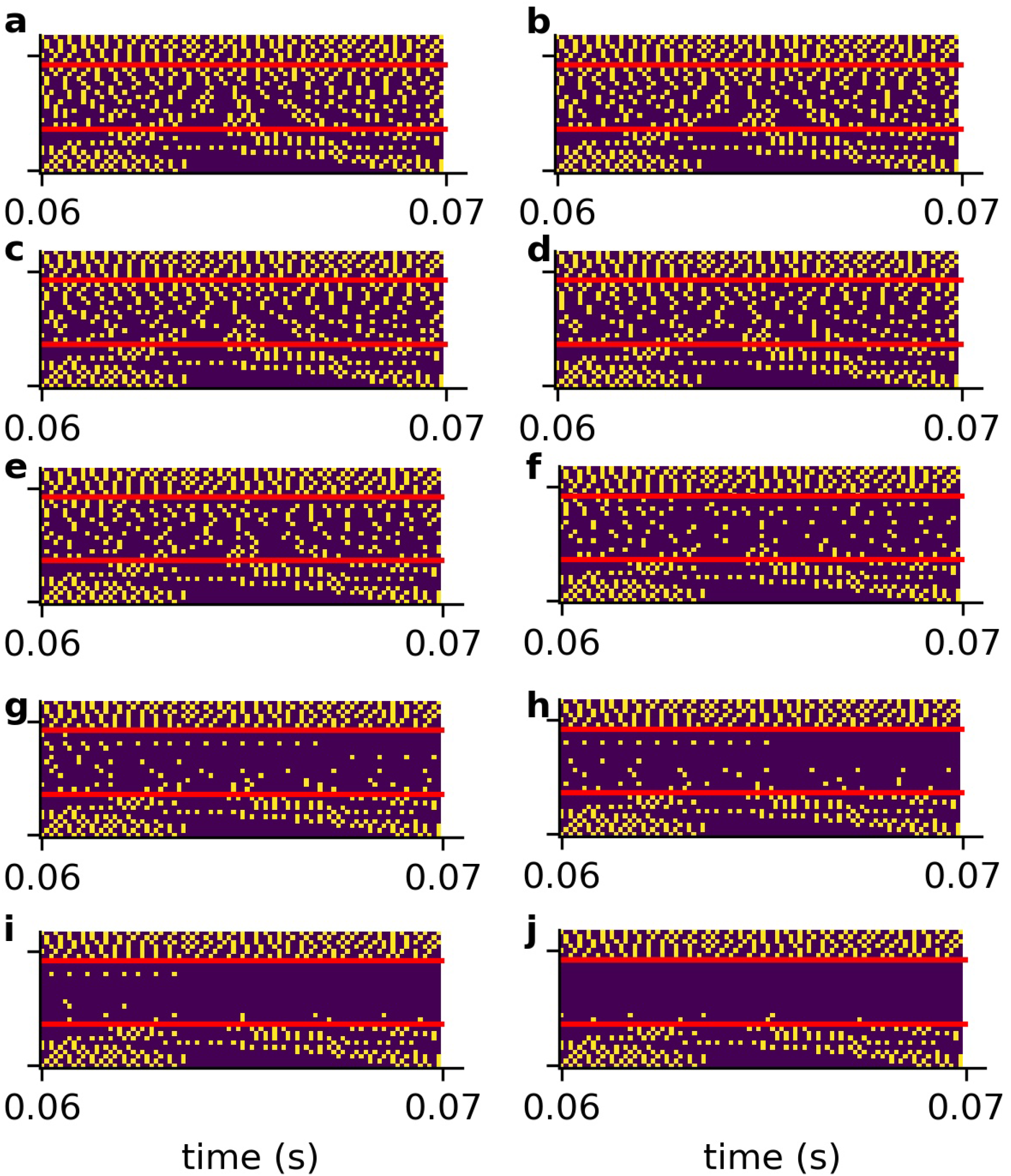
Compressed spike patterns with added hearing loss. The figure shows the down-sampled spike patterns of the same word as shown in Figure 4. The speech relevant frequency range (400Hz–4kHz) is artificially weakened (hearing loss). (a-j) refer to hearing losses 0 dB-45 dB.

### Undisturbed test data leads to a reference test accuracy of 0.37

As described above the complete auditory pathway beyond the DCN, including the superior olive, lateral lemniscus, inferior colliciculus, medial geniculate corpus, and the auditory cortex, is modeled as a deep neural network which is trained on the classification of 207 different German words (custom-made data set), or ten English words corresponding to the digits from 0 to 9 (FSDD data set [63]), respectively. In both cases the compressed, i.e. down sampled, DCN output matrices served as training and test data input.

In case of our costum-made data set, the network is exclusively trained on the data of 10 out of 12 speakers, while the remaining two speakers serve as test data. Furthermore, for network training we used only those compressed spike train matrices that correspond to the undisturbed system, i.e. without hearing loss and added noise. Due to the image-like features of the compressed spike pattern matrices, the deep neural network mainly consisted of convolutional layers. The exact architecture is shown in Figure 6a and all parameters are provided in Supplementary Table 1). For training on our costum-made data set, the test accuracy significantly decreases after 20 epochs of training (Figure 6b), and thus we applied the *early stopping procedure* [64] to prevent the network from overfitting. The trained networks were used for all further analyses with different modifications of the test data set, i.e. different hearing losses and different intensities of intrinsic noise.

**Figure 6:**
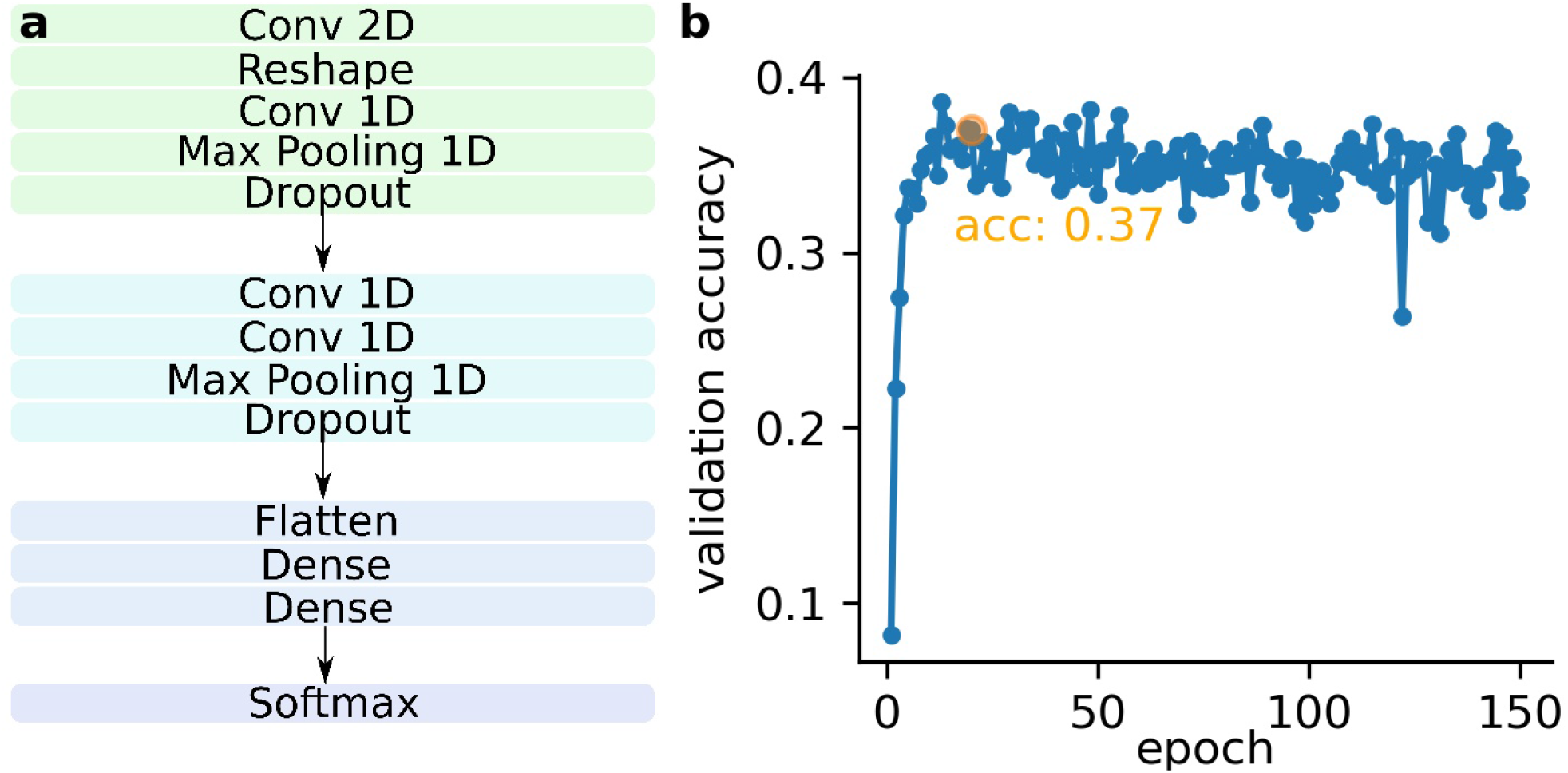
DNN architecture and validation accuracy as function of the epochs. a: Complete network architecture; The network consists of two convolutional network blocks. Each convolutional network block consists of two convolutional layers (ReLu activation), a dropout layer and a maxPooling layer. The convolutional blocks are followed by two fully connected layers (dense layers) with ReLu activation. The last layer is one further fully connected layer with softmax activation (classification layer, for exact architecture see Tab. 1). b: Validation accuracy as function of the epochs; The validation accuracy becomes worse after about 20 epochs (overfitting). The network trained for 20 epochs is used for further analysis.

### Intrinsic noise partially restores spike patterns after simulated hearing loss

To test the putative beneficial effect of intrinsic noise in case of hearing loss, we analysed spiking patterns generated with and without intrinsic noise and compared them with the corresponding undisturbed patterns (Figure 7). In Figure 7a a sample spike pattern in case of no hearing loss is shown as reference. As expected, a simulated hearing loss of 30 dB in the frequency range of 400 Hz to 4 kHz leads to a decreased spiking activity (Figure 7b), which can be partially restored by the addition of intrinsic noise with optimal intensity (Figure 7c).

**Figure 7:**
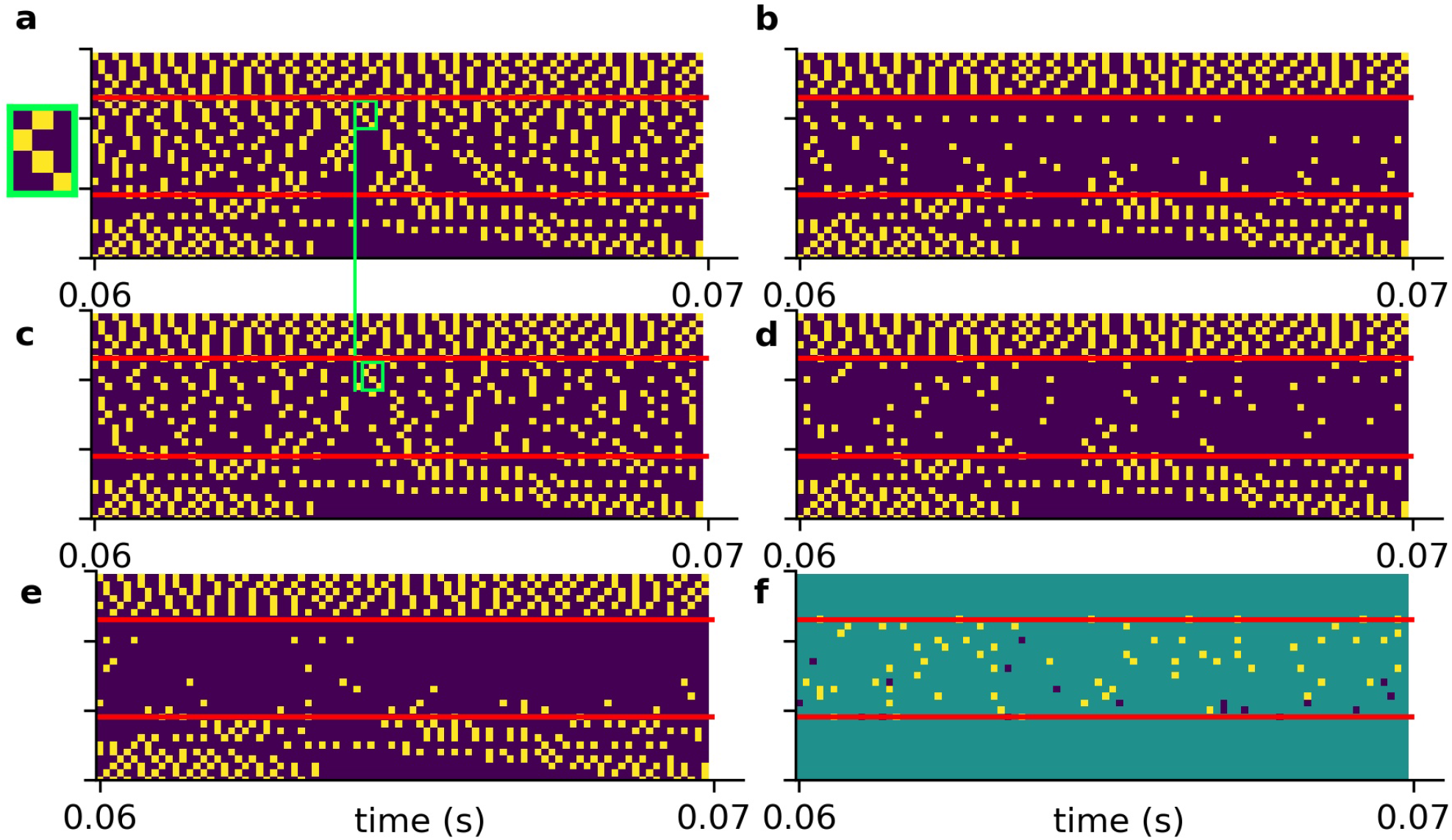
Effect of intrinsic noise on the DCN output patterns. a: Spiking without HL (same as in Figure 5a). b: Spiking with a HL of 30dB (same as Figure 5g). c: Spiking activity with HL and intrinsic noise of optimal intensity. Additional white noise increases spiking activity. d: Point-to-point comparison of spiking patterns for no HL and with HL and intrinsic noise. Shown are only spikes that occur in both cases, i.e. that are not affected by HL or that are correctly restored by noise. e: Point-to-point comparison of spiking patterns for no HL with HL and without intrinsic noise. Shown are only spikes that occur in both cases, i.e. that are not affected by HL. f: Intrinsic noise of optimal intensity not only restores spikes correctly (yellow), but also introduces false positive spikes (dark blue). Intrinsic noise restores spatio-temporal spiking patterns correctly, yet with some temporal shift (green boxes in a and c, zoom of spike pattern in green box).

A point-to-point comparison of the spikes resulting from the undisturbed system (no hearing loss) with the spikes resulting from hearing loss and additional intrinsic noise, demonstrates that there is indeed some improvement. In Figure 7d only those spikes are shown that occur in both mentioned cases. In contrast, there are less spikes resulting from hearing loss without intrinsic noise (Figure 7e). Further analysis yield that intrinsic noise not only restores spikes correctly (Figure 7f, yellow), but also introduces false positive spikes (Figure 7f, blue). However, a direct point-to-point comparison of spike patterns does not fully capture the benefit of intrinsic noise. As shown in Figure 7a, c (green boxes), intrinsic noise even restores larger spatio-temporal spiking patterns correctly, yet with some temporal shift.

### Intrinsic noise improves accuracy for speech recognition after simulated hearing loss

We also analyzed the effect of intrinsic noise on speech recognition accuracy in case of hearing loss in different scenarios. Using our costum-made data set, we investigated hearing loss in two different frequency ranges. Furthermore, using the FSDD data set, we investigated hearing loss using two different neural networks. In all cases, we find that intrinsic noise of appropriate intensity improves accuracy for speech recognition after simulated hearing loss.

### Costum-made data set and hearing loss in the frequency range of 400 Hz to 4 kHz

For the first scenario, we used a convolutional neural network (Table 1) trained on our costum-made data set. After training, we simulated a hearing loss in the frequency range of 400 Hz to 4 kHz which is known to be crucial for speech comprehension in humans [65]. The effect of improved or decreased speech comprehension is quantified by the classification accuracy of the words (test accuracy). The classification accuracy as a function of the hearing loss has a biologically plausible sigmoid shape (Figure 8a dark blue curve). The test accuracies as a function of the added noise for different hearing losses show a clear resonance curve with a global maximum (Figure 8 b). For a hearing loss of about 20 dB, the relative improvement of speech comprehension is more than doubled (Figure 8c). Furthermore, it can be shown that the optimal noise level correlates with the hearing loss (Figure 8d). This effect is plausible as for a weaker signal a higher noise amplitude is needed to lift the signal over the threshold of the LIF neurons. In summary, it can be stated that the addition of noise can lead to an improved speech comprehension for all hearing losses. This fact can be seen in Figure 8a, where the cyan curve shows the test accuracy as a function of the hearing loss with the ideal amount of added Gaussian noise.

**Table 1:** Exact paramters of the used deep convolutional network (main analysis)

| Layer | Type | input-output-dim | activation | characteristics |
| --- | --- | --- | --- | --- |
| 1 | Convolution 2D | 30, 8820, 1 ; 1, 8791, 128 | ReLu |  |
| 2 | Reshape | 1, 8791, 128; 8791, 128 |  |  |
| 3 | Convolution 1D | 8791, 128; 8782, 128 | ReLu |  |
| 4 | MaxPooling 1D | 8782, 128; 4391, 128 |  | poolsize: 2 |
| 5 | DropOut | 4391, 128; 4391, 128 |  | droupout: 0.5 |
| 6 | Convolution 1D | 4391, 128; 4391, 128 | ReLu |  |
| 7 | Convolution 1D | 4391, 128; 4390, 128 | ReLu |  |
| 8 | MaxPooling 1D | 4390, 128; 2195, 128 |  | poolsize: 2 |
| 9 | DropOut | 2195, 128; 2195, 128 |  | droupout: 0.5 |
| 10 | Flatten | 2195, 128; 280960 |  |  |
| 11 | Dense | 280960; 150 | ReLu |  |
| 12 | Dense | 150; 50 | ReLu |  |
| 13 | Dense | 50; 207 | Softmax |  |

**Figure 8:**
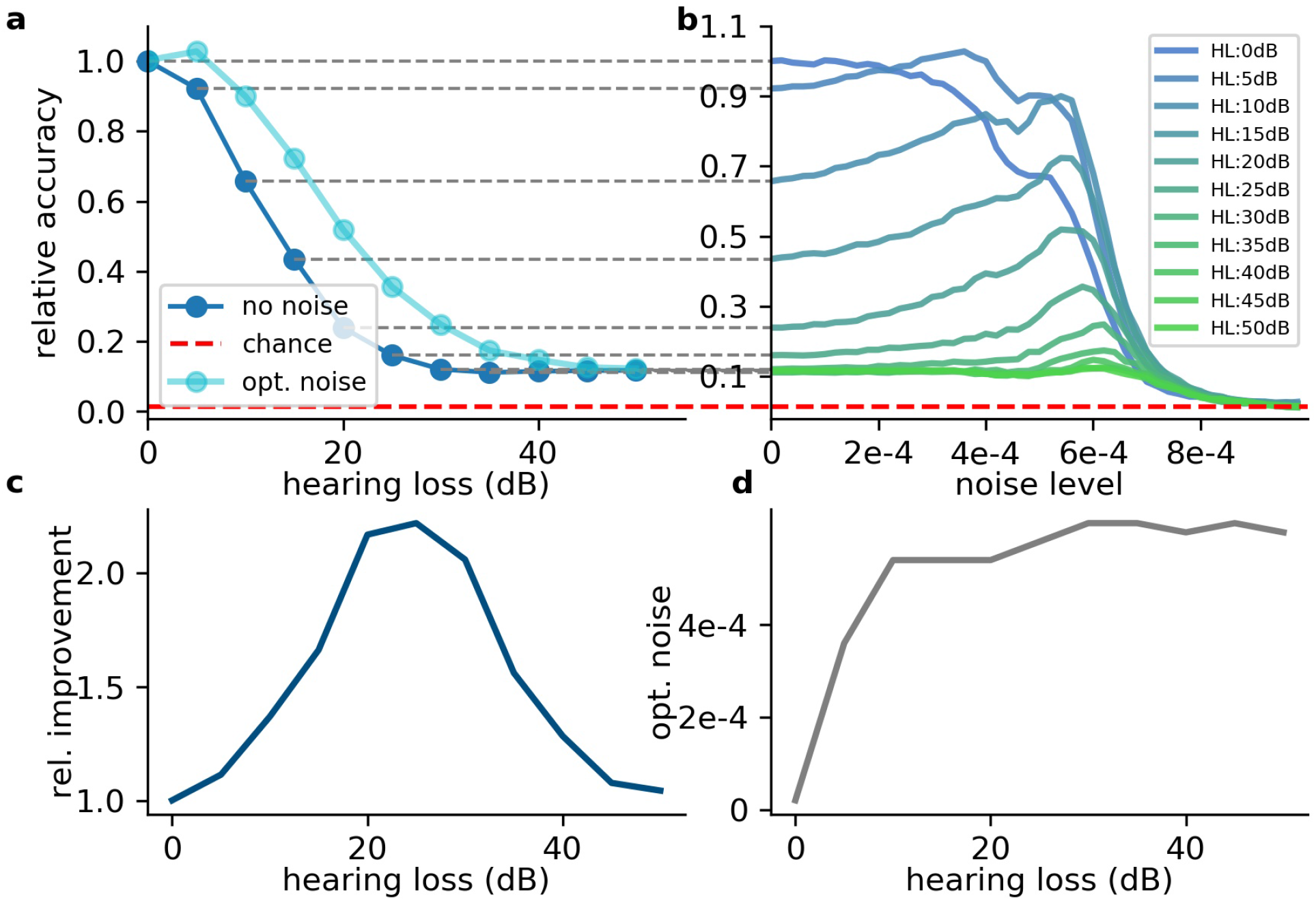
Effect of SR on speech recognition. a: The curve shows the relative accuracy of the trained neural network as a function of the hearing loss (red dashed line: chance level; 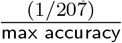). The hearing loss (5–50 dB, 5 dB steps, frequency range of HL: 400 Hz–4000 Hz) was implemented in the test data set and propagated through the pre-trained network. Thus, the cochlea output was multiplied with an attenuation factor 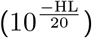. This output was then transformed using the integrate-and-fire neurons and fed in the neural network. b: Relative accuracy as a function of the applied noise level for different hearing losses. Resonance curves with one global maximum at a certain noise level >0 could be shown. c: Best relative improvement as a function of the hearing loss. d: Optimal noise level as a function of the hearing loss

### Costum-made data set and hearing loss in the frequency range above 4 kHz

Since many people suffer from hearing losses in the high frequency range [66]. In the next step, the stochastic resonance effect is analyzed for a high frequency range hearing loss starting at a frequency of 4 kHz. It can be shown that the high frequency loss does not affect the speech comprehension abilities in the same manner as hearing losses in the critical frequency range between 400 Hz and 4 kHz (Figure 9a). The relative accuracy does not drop below a value of 50%. Thus, the effect of stochastic resonance is also reduced (Figure 9b), which means a maximal relative improvement of approximately 10% (Figure 9c, d). Furthermore, there is no real resonance curve with one maximum at a certain noise frequency but a second maximum at a higher noise level (Figure 9b). To put it in nutshell, we can state that the addition of noise can lead to a significant improvement of speech comprehension.

**Figure 9:**
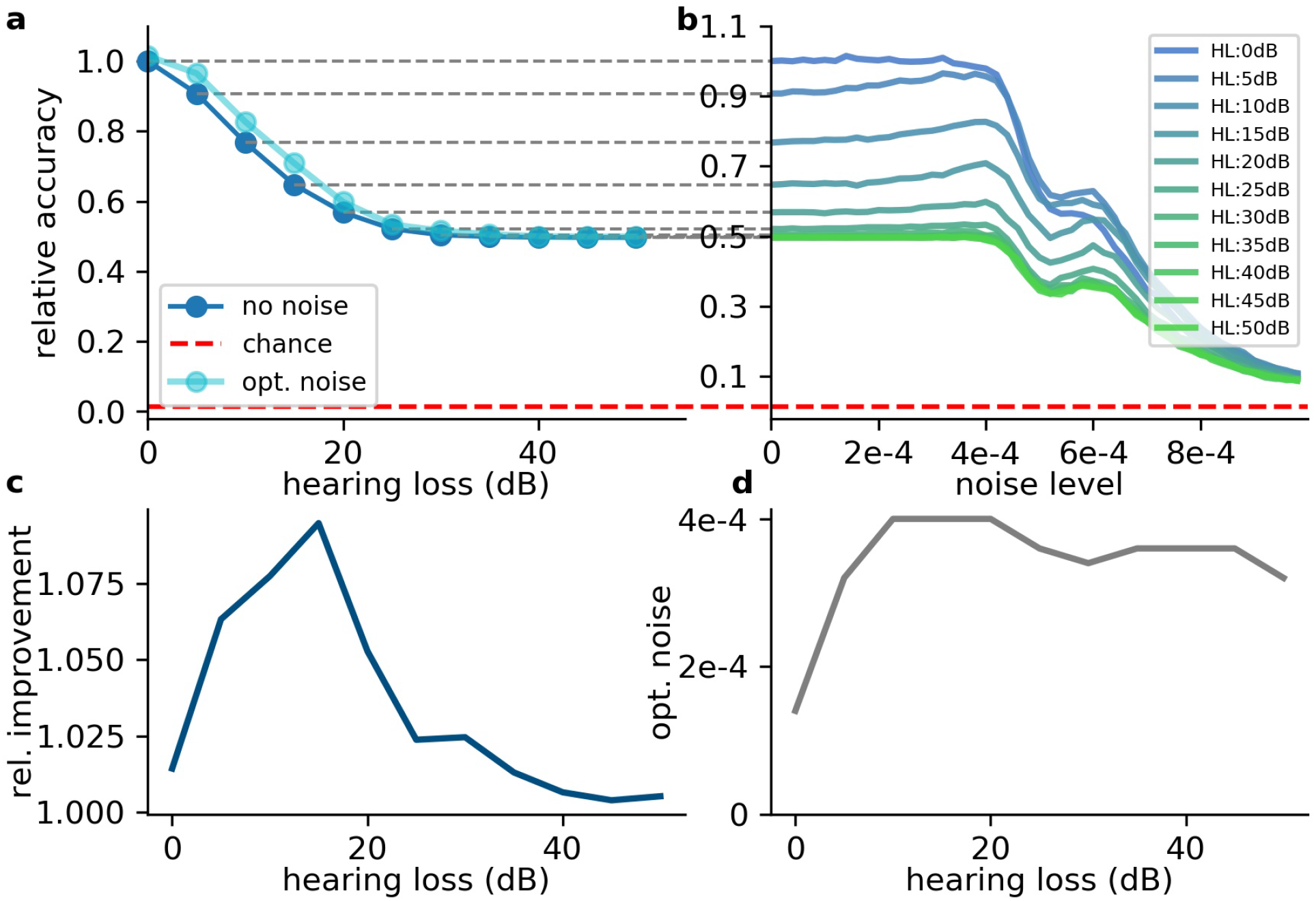
Effect of SR on speech recognition (high frequency hearing loss) Same analysis as shown in Figure 8 for high frequency hearing loss. a: The plots show the relative accuracy of the trained neural network as a function of the hearing loss (red dashed line: chance level). The high frequency hearing loss lead to different effects(10–50 dB, 10 dB steps, frequency range of HL: above 4000 Hz). b: The relative accuracy as a function of the noise has no clear maximum above the value for no added noise (nearly no SR). Furthermore, a second local maximum occurs. c: The best relative improvement does not significantly increase over 10%. d: Optimal noise level as a function of hearing loss shows similar behavior as for the hearing loss in the speech relevant frequency range (cf. Figure 8).

### FSDD data set and hearing loss in the frequency range above 400 Hz

In order to demonstrate that this effect is not limited to a certain data set, language or neural network architecture, we repeated our analyses using two further neural networks, a convolutional neural network (Supplementary Table 2) and a network with Long-Short-Term-Memories (Supplementary Table 3), both trained and tested with the FSDD data set (Figure 10). A hearing loss in the critical frequency range for speech comprehension leads to a decrease in the classification accuracy (10a for the convolutional network and 10c for the Long-Short-Term-Memory network). Furthermore, the stochastic resonance effect in terms of a clear resonance curve with one maximum can be observed (Figure 10b, d).

**Figure 10:**
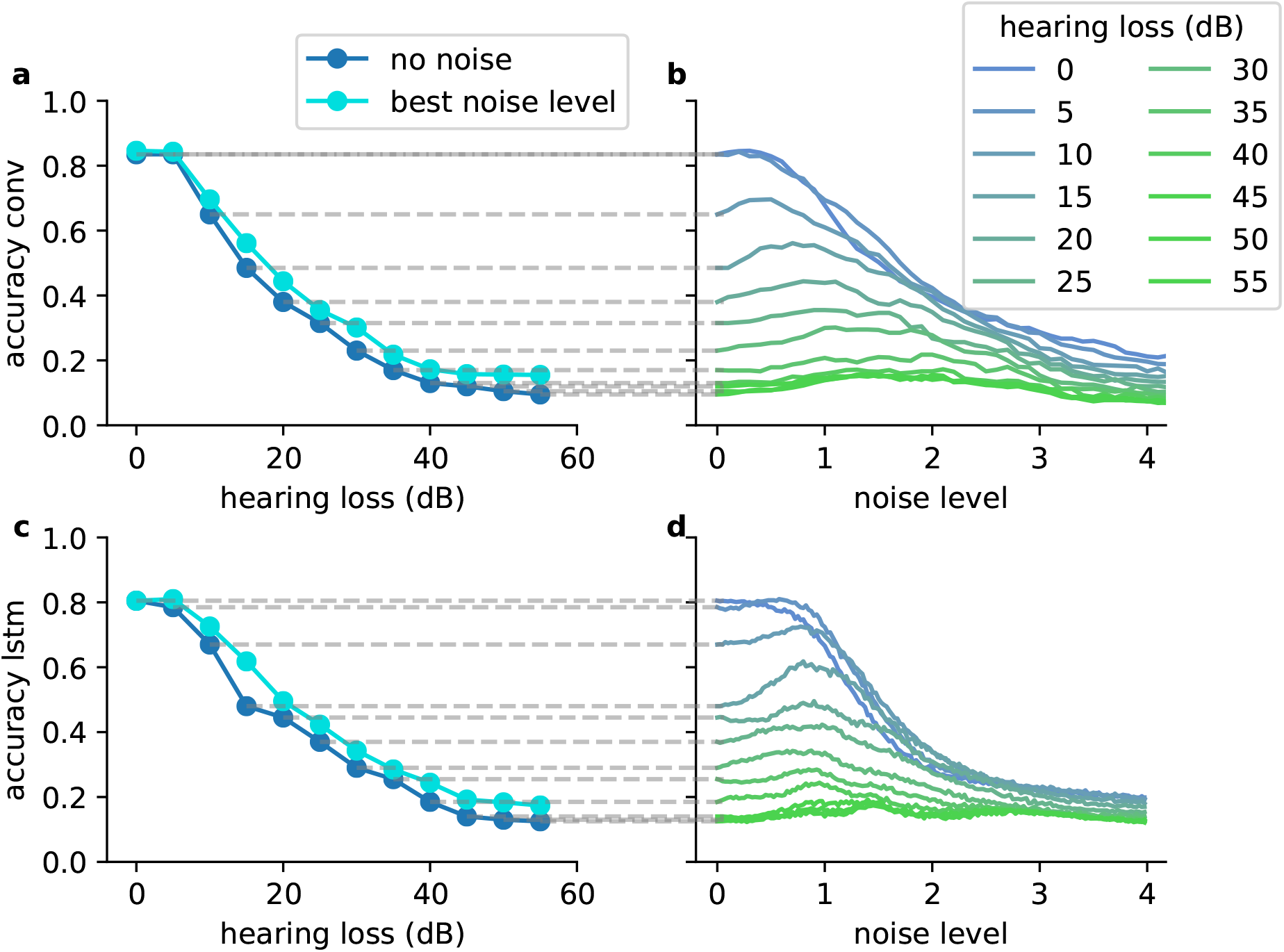
The SR resonance effect in different network architectures using the FSDD data set. a: The plot shows the test accuracy as a function of the applied hearing loss for a deep convolutional network architecture (dark blue, starting at 400 Hz, exact network architecture shown in Table 2) trained on English words (digits: 0-9). The impaired speech comprehension by the hearing loss can be partly compensated by adding Gaussian noise (stochastic resonance). The cyan curve shows the improvement of speech comprehension for the optimal noise level (maxima values in b). b: Test accuracy for different hearing losses (shades of blue) as a function of the added noise. The maxima show that SR can help to restore speech comprehension. c: Similar analysis as shown in (a) for a two layer LSTM network (exact network architecture shown in Table 3); d: Similar analysis as shown in (b) for the LSTM architecture. The improvement of speech perception in impaired systems (hearing loss) is a universal principle and does not depend on the used neural network.

**Table 2:** Exact paramters of the used deep convolutional network (FSDD data set)

| Layer | Type | input-output-dim | activation | characteristics |
| --- | --- | --- | --- | --- |
| 1 | Convolution 2D | 9131, 30, 1; 9102, 1, 32 | relu |  |
| 2 | MaxPooling 2D | 9102, 1, 32; 4551, 1, 32 |  | poolsize: (2, 1) |
| 3 | DropOut | 4551, 1, 32; 4551, 1, 32 |  | droupout: 0.2 |
| 4 | Convolution 2D | 4551, 1, 32; 4547, 1, 64 | relu |  |
| 5 | MaxPooling 2D | 4547, 1, 64; 2273, 1, 64 |  | poolsize: (2, 1) |
| 6 | DropOut | 2273, 1, 64; 2273, 1, 64 |  | droupout: 0.2 |
| 7 | Convolution 2D | 2273, 1, 64; 2272, 1, 32 | relu |  |
| 8 | MaxPooling 2D | 2272, 1, 32; 1136, 1, 32 |  | poolsize: (2, 1) |
| 9 | DropOut | 1136, 1, 32; 1136, 1, 32 |  | droupout: 0.2 |
| 10 | Flatten | 1136, 1, 32; 36352 |  |  |
| 11 | Dense | 36352; 400 | relu |  |
| 12 | DropOut | 400; 400 |  | droupout: 0.2 |
| 13 | Dense | 400; 50 | relu |  |
| 14 | Dense | 50; 10 | softmax |  |

**Table 3:** Exact paramters of the used LSTM network (FSDD data set)

| Layer | Type | input-output-dim | activation | characteristics |
| --- | --- | --- | --- | --- |
| 1 | GroupToBatches | 9000, 30; 45, 6000 |  |  |
| 2 | LSTM | 45, 6000; 45, 200 | tanh |  |
| 3 | DropOut | 45, 200; 45, 200 |  | droupout: 0.5 |
| 4 | LSTM | 45, 200; 45, 100 | tanh |  |
| 5 | DropOut | 45, 100; 45, 100 |  | droupout: 0.5 |
| 6 | TimeDistributed Dense | 45, 100; 45, 100 |  |  |
| 7 | DropOut | 45, 100; 45, 100 |  | droupout: 0.5 |
| 8 | TimeDistributed Dense | 45, 100; 45, 10 |  |  |

## Discussion

In this study, we demonstrated with a computational model of the auditory system that noise added to the DCN may improve speech recognition after hearing loss, by means of SR. The relative benefit of SR turned out to be largest for hearing losses between 20 and 30 dB.

Because SR works by partly restoring missing information in the input data, adding noise improves the classification accuracy of the neural network even after the training period is finished. This stands in contrast to machine learning techniques that achieve an increased robustness and generalization ability by purposefully using noisy training data from the beginning [67], or by adding artificial noise during the training period [68, 69].

In our work, we first train the neural network for speech recognition, then simulate a hearing loss, and finally reduce this loss by adding noise. This approach is biologically plausible, as also the brain is trained on speech recognition during childhood [70, 71], where hearing ability is usually optimal (Indeed, hearing impairment in childhood can lead to problems in language acquisition, which cannot be fully cured in adulthood [72]). In the coarse of a lifetime, hearing ability becomes permanently [73, 74] or temporary worse [75], often due to high amplitude sound exposure.

We have proposed that hearing ability can be restored by a control cycle embedded in the brainstem, along the auditory pathway, which uses internal neural noise to exploit the effect of stochastic resonance [22]. Thus, it is supposed that the neural activity in damaged frequency channels is up-regulated by internally generated noise to restore hearing within this frequency range.

Overshooting of this noise up-regulation is proposed to be the origin of tinnitus [22]. Our model could provide an interesting explanation for overshooting internal noise: In our simulation of high frequency hearing loss, we found that the accuracy as a function of the added noise has not only a single maximum, as expected for a resonance curve, but features a second maximum at a higher noise level (Figure 9b). If the neural control cycle would be drawn to this secondary maximum, this might explain an overshooting of the neural noise and the corresponding emergence of tinnitus [21, 22].

Another potential cause of tinnitus arises from the fact that phase locking, the encoding of a signal’s phase information in neural spike trains, is only possible for frequencies up to 4 kHz, the maximum spike rate of the DCN neurons (Figure 3a).

The stochastic resonance effect probably works only below this limit frequency, and thus it is not clear whether (or how) the neural control system compensates for the hearing loss in the frequency range above 4 kHz, as it has no real maximum to optimize for. Potentially, the tuning of the noise parameters in this frequency regime is done only by random trial. This model would fit to the observation that tinnitus mainly occurs in the high frequency range [23].

We were able to show that neural noise could potentially help to increase speech comprehension in neural systems in a computational model of the auditory pathway. An improvement of up to a factor of 2 is possible. This model provides new insights how the auditory system optimizes speech comprehension on small time scales. Furthermore, we could give a mechanistic explanation of the development and characteristics of tinnitus perception. These finding could have a major impact on medical treatment of phantom perceptions, but on the other hand raises new research questions in the field of engineering. It would be interesting to implement these principles to improve artificial systems trained on speech comprehension. Furthermore, the effect of stochastic resonance could be used to improve sensory systems [21].

Our study provides evidence that an interplay of machine learning and neuroscience helps on the one hand to raise understanding of the function of biological neural networks (e.g [3, 76, 77]), an emerging science strand referred to ”cognitive computational neuroscience” [78]. On the other hand, basic principles from nature –such as stochastic resonance– can be derived to improve artificial neural systems, which is called ”machine behavior” [79].

## Methods

### Computational Resources

The simulations were run on a desktop computer equipped with an i9 extreme processor (Intel) with 10 calculation cores. Furthermore, the machine learning was run on the same computer on two Nvidia Titan XP graphical processor units. To test the validity of our calculations the simulations were performed on two different code bases. The main results based on our own speech data set are mainly based on Numpy [80] and SciPy [81] calculations. The convolutional network was implemented in Keras [82] with Tensorflow [83] back-end. All main results were confirmed by analyzing a standard speech data set–the so called Jakobovski free spokeb digit data set (FSDD) [63], containing spoken numbers from 0 to 9 in English language in accordance to the MNIST data set with written digits in this range [84]. This was done using a completely new code base exclusively build of KERAS layers. Thus, a custom-made KERAS layer implemented as sinc FIR filters for the cochlea layer as well as the leaky-integrate-and-fire neurons were implemented. All plots were created using the Matplotlib Python library [85] and plots were arranged using the pylustrator [86].

### Cochlea model

The cochlea is simulated as 30 butterworth bandpass filters (3rd order) with no overlapping bands. These 30 bandpass filters are a simplification of the more than 3000 inner hair cells of the human cochlea [87]. In contrast to other complex cochlea models [88,89], this simplification of the dynamics of the inner hair cells was chosen to derive basic principles and to increase interpretability. The center frequencies (of the bandpass filters) are between 100 Hz (minfreq.) and 10 kHz (maxfreq.) including the complete frequency range needed for speech comprehension. The center frequencies are chosen to grow exponentially (centerfreq. = minfreq. · factor*^i^* with *i* ∈ {0,1,…,29} and 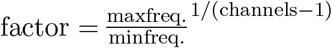). Thus, for higher frequencies the spacing of the center frequencies becomes larger in analogy to the tonotopy of the human cochlea [61,65]. The width of the bandpass filters is defined as [centerfreq. · factor^−0.5^, centerfreq. · factor^0.5^].

### Dorsal cochlear nucleus model

The dorsal cochlear nucleus (DCN) was modeled as 30 leaky integrate-and-fire (LIF) neurons [60], each of these neurons is connected to one frequency channel of the cochlea. Thus, no lateral inhibition was realized to focus on the core effects. The maximum spiking rate of the simulated LIF neurons is approximately 4 kHz (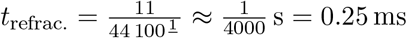, *t*_refrac_.: refractory time), which is much s more than the maximum spiking rate of a biological neuron (250 Hz) [90]. Thus, in the simulation one LIF neuron represent approximately 16 real neurons. The recruitment of several neurons to increase the frequency range in which phase coupling is possible is a core concept within the dorsal cochlear nucleus [91]. The numerical integration of the LIF neurons was performed using the ”Euler” method, as this method lead to the lowest computational complexity compared to ”Heun” and ”Runge Kutte”—being standard integration techniques [92]—without causing significant inaccuracies.

### Brain stem and cortex model

The neural processing stages of the auditory pathway above the DCN including superior olive, lateral lemniscus, inferior colliculus, medial corpus geniculatum in the thalamus, and auditory cortex are modeled as a deep neural network [61]. For our main simulations we used a Deep Convolutional Neural Network [93] to gain independence from translations of the data (for architecture see Fig. 6a, for exact parameters cf. Supplements Tab. 1). Furthermore, we used also Deep LSTM networks [94] to double-check the validity and universality of the beneficial effects of intrinsic noise (Supplements Table 3).

### Data sets for speech recognition

#### Custom-made data set

Our custom-made data set was recorded from 12 different speakers (6 male, 6 female) in a range of 20–61 years. The data was recorded with a sampling rate of 44.1 kHz bit using Audacity. Each participant had to speak the 207 most common German words 10 times each. After recording the data was labeled using forced alignment and cut into 1 s intervals. The data from 10 participants served as training data set, whereas the data from the two other speakers was used as test data set. All evaluations, i.e. simulated hearing loss and effect of intrinsic noise, were based on the modified test data.

#### FSDD data set

The second used data set is an open data set consisting of spoken digits (0-9) –in analogy to the MNIST data set– in English. The data set is sampled with 8 kHz and consists of 2000 recorded digits from 4 speakers [63]. Here the first five repetitions of for each speaker and each digit are used as test data, the respective remaining 45 repetitions serve as training data.

## Acknowledgments

This work was funded by the Deutsche Forschungsgemeinschaft (DFG, German Research Foundation): grant KR5148/2-1 to PK – project number 436456810, and the Emergent Talents Initiative (ETI) of the University Erlangen-Nuremberg (grant 2019/2-Phil-01 to PK), and the Interdisciplinary Center for Clinical Research (IZKF) at the University Hospital of the University Erlangen-Nuremberg (grant ELAN-17-12-27-1-Schilling to AS).

The authors are grateful for the donation of two Titan Xp GPUs by the NVIDIA Corporation.

We thank Martin Haller for technical assistance, and Holger Schulze for providing us access to the lab.

Finally, we wish to thank the twelve speakers who lend us their voices for generating the costum-made speech data set.

## Author contributions

PK and AS designed the study. AZ created the costum-made data set. AS, RG and PK implemented the model. AS, RG, CM, AM and PK discussed the results. AS, PK, RG, CM and AM wrote the manuscript.

